# Network-targeted TMS modulates task-related striatal activity during motor skill learning

**DOI:** 10.64898/2026.03.17.712285

**Authors:** Sungbeen Park, Junghyun Kim, Yonghyun Kwon, Sungshin Kim

## Abstract

The striatum, a critical hub for motor skill learning, is located deep within the subcortical region, making noninvasive stimulation particularly challenging. Nevertheless, recent studies suggest that transcranial magnetic stimulation (TMS) can modulate subcortical activity indirectly by targeting functionally connected cortical areas. In this study, we applied TMS to the dorsolateral prefrontal cortex (DLPFC) immediately before the fMRI session measuring task-related activity in the striatum during motor learning. We examined whether continuous theta-burst stimulation (cTBS) and high-frequency stimulation (20 Hz) could modulate motor learning and associated striatal responses with opposing effects. There was no significant effect of either stimulation condition on the overall motor learning performance. However, cTBS significantly reduced performance-related striatal activity, while 20 Hz stimulation did not show any modulatory effect. These findings demonstrate that cTBS targeting the corticostriatal network can suppress striatal activity and suggest its potential use in clinical trials for treating disorders such as addiction associated with hyperactive striatal responses.

## Introduction

Motor skill learning in humans emerges from dynamic interactions between cortical and subcortical networks, with the dorsolateral prefrontal cortex (DLPFC), primary motor cortex (M1), and striatum forming key nodes within these systems. The DLPFC supports cognitive control over goal-directed actions [1], M1 implements motor output and experience-dependent plasticity, and the striatum integrates inputs from these and other regions to link motor planning with execution within cortico-striatal circuits [2–4]. Anatomical and functional evidence indicates that anterior and posterior subdivisions of the striatum form partially segregated but parallel loops with distinct frontoparietal territories, establishing anterior “cognitive” and posterior “sensorimotor” pathways that differentially contribute across learning stages [4–6]. Early in skill acquisition, when action selection must remain flexible, prefrontal–anterior striatal circuits are preferentially engaged, whereas later automatized and habitual performance increasingly relies on parietal/sensorimotor–posterior striatal loops [7–11]. Converging human and animal work suggests that this anterior-to-posterior shift reflects a conserved organizational principle whereby cognitive and sensorimotor mechanisms interact hierarchically over time during motor skill learning [12,13].

Despite substantial progress in characterizing these cortico-striatal dynamics, their causal contributions to human motor learning remain incompletely understood. Noninvasive brain stimulation, particularly transcranial magnetic stimulation (TMS), offers a powerful approach for testing such causal roles by introducing transient perturbations at specific cortical nodes and probing resulting changes in behavior and network interactions [14,15]. Studies applying TMS after practice have shown that stimulation can differentially modulate motor memory consolidation depending on the structure of practice, highlighting the potential of TMS-based interventions to selectively engage and shape specific neural circuits [16]. At the neurochemical level, positron emission tomography (PET) work demonstrates that repetitive TMS (rTMS) over the DLPFC increases dopamine release in the caudate nucleus, whereas rTMS over M1 induces dopamine release in the dorsolateral putamen [17,18]. These findings indicate that cortical stimulation can influence fronto-striatal communication at multiple levels and provide a mechanistic basis for TMS-induced modulation of learning-related plasticity.

Neuroimaging studies further underscore that motor skill learning involves coordinated plastic changes across motor cortex and striatum, with the putamen playing a central role in sequence learning tasks such as finger tapping and being sensitive to inhibitory rTMS applied to M1 [19–21]. More recent work shows that DLPFC stimulation modulates task-dependent functional connectivity within cortico-striatal networks and impacts memory consolidation as well as large-scale systems integration [22–26]. Our previous studies using de novo motor skill learning paradigms similarly revealed robust task-evoked activity not only in sensorimotor regions but also in higher-order cognitive areas, including the ventromedial prefrontal cortex (vmPFC) and the striatum, supporting integrative models in which multiple memory systems jointly contribute to performance [8,27]. Collectively, these findings motivate a network-level framework in which TMS can be used to shape the coordinated engagement of distinct cortico-striatal loops underlying motor learning [28–32].

However, it remains unclear whether TMS can modulate task-evoked striatal activity during the acquisition of more complex motor skills that impose higher cognitive demands than simple motor tasks. The present study addresses this gap by examining how TMS applied to the DLPFC modulates striatal activation indirectly, via fronto-striatal network targeting, during a complex de novo motor skill learning task. To this end, we employed continuous theta-burst stimulation (cTBS) and high-frequency rTMS (20 Hz) as complementary protocols expected to exert opposing effects on motor performance and task-related striatal activity [15,33]. We hypothesized that these two stimulation paradigms would bidirectionally modulate motor skill acquisition and striatal responses through their network-level impact [33–35]. Building on evidence that anterior striatal regions such as the caudate support early stages of novel motor sequence learning, whereas posterior regions such as the putamen support automatization [4,7,8,11], we further tested whether cTBS applied to the DLPFC versus M1 differentially regulates these striatal subregions. Specifically, we predicted that inhibitory stimulation of the DLPFC would suppress task-related activity in the caudate, whereas excitatory stimulation would enhance caudate responses. In adㅊdition, because inhibitory stimulation of M1 is expected to exert more pronounced effects during later learning phases, we used M1 cTBS as a comparison condition to DLPFC stimulation during early learning. This experimental design aims to provide causal evidence for distinct contributions of anterior and posterior cortico-striatal circuits across different stages of complex motor skill learning.

## Materials and Methods

### Participants

Seventy-one healthy young adults were recruited, of whom 50 (27 females, 23 males; mean age = 23.1 years, SD = 3.2 years) completed all behavioral and fMRI experiments on Day 1, as well as TMS and fMRI experiments on day 2. All 50 participants met the health criteria for fMRI and TMS (i.e., no history of neurological disorders, claustrophobia, extreme noise sensitivity, color blindness, or other contraindications; normal or corrected-to-normal vision). In addition, they were all right-handed according to the modified version of the Edinburgh Handedness Inventory [36]. Data of 21 participants were excluded, because 16 participants did not come to the experiment place without telling a reason or without a specific reason; two participants refused TMS session; one participant had undetachable earrings; data of one participant was lost due to technical reasons; and one participant never reached the first column of the 5×5 grid.

Participants were randomly assigned into 3 groups: continuous theta-burst stimulation (cTBS) on M1, cTBS on DLPFC, and 20-Hz repetitive TMS (rTMS) on DLPFC. Of the 50 participants, 17 participants (9 females, 8 males; mean age = 23.8 years, SD = 4.3 years) were assigned into the M1-cTBS group, 17 (9 females, 8 males; mean age = 22.8 years, SD = 2.5 years) were in the DLPFC-cTBS group, and the other 16 (9 females: 7 males; mean age = 22.8 years, SD = 2.4 years) were in the DLPFC-20Hz group. For a baseline condition without TMS, we included data of 30 participants from our previous study using the same task [8], in which the first three runs (“trained”) performed during participants’ initial visit were used as the baseline.

The mean interval between the two visits was 11.8 ± 14.4 days (range: 1–47 days). They provided an informed consent before the Day-1 experiment and received a monetary reward for participation. The entire experimental procedure adhered to the Declaration of Helsinki and was approved by Sungkyunkwan University Institutional Review Board (IRB No. 2018-05-003-032).

## Experimental procedures

### Day 1

#### Calibration

During the initial visit, participants completed a calibration procedure utilizing an MR-compatible data glove (14 Ultra, 5DT Technologies), consistent with established behavioral paradigms employing this device [37]. The glove was equipped with 14 sensors: two per finger to measure flexion excluding proximal joints and four sensors dedicated to capturing finger abduction. Over a two-minute calibration phase, participants freely flexed and extended the fingers of their right hand without any constraints. Sensor outputs were sampled as scalar values at 60 Hz. The resulting dataset’s covariance matrix was subjected to principal component analysis (PCA), from which the first two principal components (PC1 and PC2) were extracted to form the linear mapping matrix A (i.e., the PC1 corresponded to motion along the x-axis and the second PC to motion along the y-axis). The offset p_0_ was computed to ensure that the mean hand posture positioned the cursor at the center of the display screen. These procedures were conducted using MATLAB version 2019a. Consequently, during the main task, the 14-dimensional sensor output vector ℎ obtained from the glove was mapped to the two-dimensional cursor position p on the monitor according to the linear transformation (i.e., p = Aℎ + p_0_):

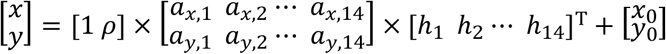

where *ρ* denotes the relative ratio of the eigenvalues associated with PC1 and PC2. Because the PC1 always accounts for at least as much variance as the PC2, the components were adjusted by the square root of their respective eigenvalues to equalize the challenge across the x- and y-directions.

#### Familiarization

Participants first engaged in a familiarization phase prior to training, attempting to reach all 25 targets arranged in the 5×5 grid. This procedure ensured that the entire workspace was accessible. All participants reached the targets successfully, and no modification of the original mapping matrix was necessary.

#### Resting-state fMRI session

Participants underwent an 8-min and 30-s resting-state fMRI session while wearing the data glove on their right hand, in preparation for the subsequent localizer session. They were instructed to lie comfortably, relax, and avoid specific thoughts or body movements. Foam padding minimized head motion. Resting-state data were not analyzed.

#### Localizer session

A localizer session identified regions associated with random finger movements. Participants positioned their right hand comfortably and moved their fingers freely at a natural pace during “Move” cues and rested during “Stop” cues (1 min each). Participants could not view their hands throughout the fMRI experiment. Four Move-Stop pairs were presented (total: 8 min). Localizer data were not analyzed.

#### T1 image session

High-resolution T1-weighted anatomical images were acquired, completing the Day-1 protocol. These images were used to determine individualized TMS target sites for each participant.

#### Identification of TMS targets

TMS target sites were defined using standardized Montreal Neurological Institute (MNI) coordinates. The primary motor cortex (M1) target was set at MNI coordinates [-38, -22, 52], corresponding to the “hand-knob” area [38]. The dorsolateral prefrontal cortex (DLPFC) target was determined using coordinates reported in a previous study; the Talairach coordinates provided in that work were converted to MNI space, yielding MNI coordinates of [-42, 32, 32] [18]. For each participant, a T1-weighted anatomical image acquired on the Day-1 experiment was preprocessed for spatial normalization. An affine transformation matrix was estimated between each participant’s original anatomical space and the MNI standard space, and this matrix was then applied to the MNI-defined target coordinates to identify the corresponding stimulation targets in each individual’s original anatomical space.

### Day 2

Participants returned for the Day-2 experiment. To enable immediate fMRI scanning following TMS, participants were prepared for the Day-2 fMRI session prior to stimulation. The preparation included a safety check for MRI compatibility and a brief task explanation: participants would interact with a 5×5 grid, reaching targets by moving their right fingers.

#### TMS Procedures

The TMS procedure was conducted using an ALTMS machine (Remed Co Ltd, Korea) with a figure-of-eight coil of 200 mm (width) x 100 mm (height) x 45 mm (thickness). The stimulation targets, M1 and DLPFC, were localized via a neuro-navigation system (Brainsight, Rogue Research, Montreal, Canada) integrated with a Polaris camera. Prior to stimulation, participants were seated in a TMS chair, and four anatomical fiducial points, the nasion, tip of the nose, and left and right preauricular points, were digitized to establish head mesh alignment. Individual target coordinates, determined during the Day-1 session, were subsequently imported into the system. During online calibration, spatial correspondence between each participant’s brain anatomy and the virtual brain model displayed on the monitor was confirmed, after which the TMS coil was calibrated to ensure accurate tracking of its center using the neuro-navigation system.

Stimulation intensity was determined individually based on the resting motor threshold (RMT). Participants were instructed to rest their forearms on the chair armrests with palms facing upward and fingers relaxed. Single-pulse TMS was applied over the primary motor cortex (M1) with the coil positioned tangentially to the scalp. RMT was defined as the lowest stimulation intensity that elicited a visible motor response in the contralateral hand in at least five out of ten consecutive trials. Due to practical constraints, EMG recording was not feasible; therefore, RMT was assessed by visual observation of motor twitches, a method previously employed in TMS research. Although visual inspection may slightly overestimate RMT compared to EMG-based procedures, the same criterion was consistently applied across all participants. Statistical analysis confirmed no significant differences in RMT between groups (F(2,47) = 0.315, p = 0.731), indicating that the chosen method did not introduce systematic bias in group comparisons.

Following RMT determination, either continuous theta-burst stimulation (cTBS, 40 s) or repetitive TMS at 20 Hz (20 min) was applied at 80% of RMT. The cTBS protocol consisted of triplet pulses at 50 Hz repeated every 200 ms for 40 s. The 20 Hz rTMS protocol consisted of 40 pulses delivered within 2 s, repeated every 30 s for a total duration of 20 min, following previously established procedures [33–35]. During stimulation, coil position was maintained with real-time feedback from Brainsight, and a coil holder was employed to stabilize the coil throughout the session.

#### Main fMRI experiment

Immediately following TMS, participants were escorted to the MRI suite, positioned supine on the scanner bed with foam pads to minimize head motion. An MR-compatible data glove was fitted to their right hand. Visual task feedback was presented on a monitor viewed via mirror, preventing direct observation of their hand. A central fixation cross was displayed throughout.The task was identical to that used in our previous study (Choi et al. 2020); it is briefly described the task (see Supplementary Movie S1 for demonstrations).

The experiment comprised three runs, each consisting of 97 trials, totaling 291 trials. Each trial lasted 5 s and was presented consecutively without inter-trial intervals, resulting in a run duration of approximately 8 minutes. At the start of each run, a 5×5 grid was displayed on the screen, with the first target cell (target 1) shaded gray, and a cross-shaped cursor (“+”) positioned at the center of the screen (Fig 1D). Participants manipulated their right-hand fingers to move the cursor into the target cell, which turned red upon entry, providing immediate feedback. They were instructed to hold the cursor in the target as long as possible, sustaining the corresponding finger configuration (calibrated on Day 1).

**Fig 1.**
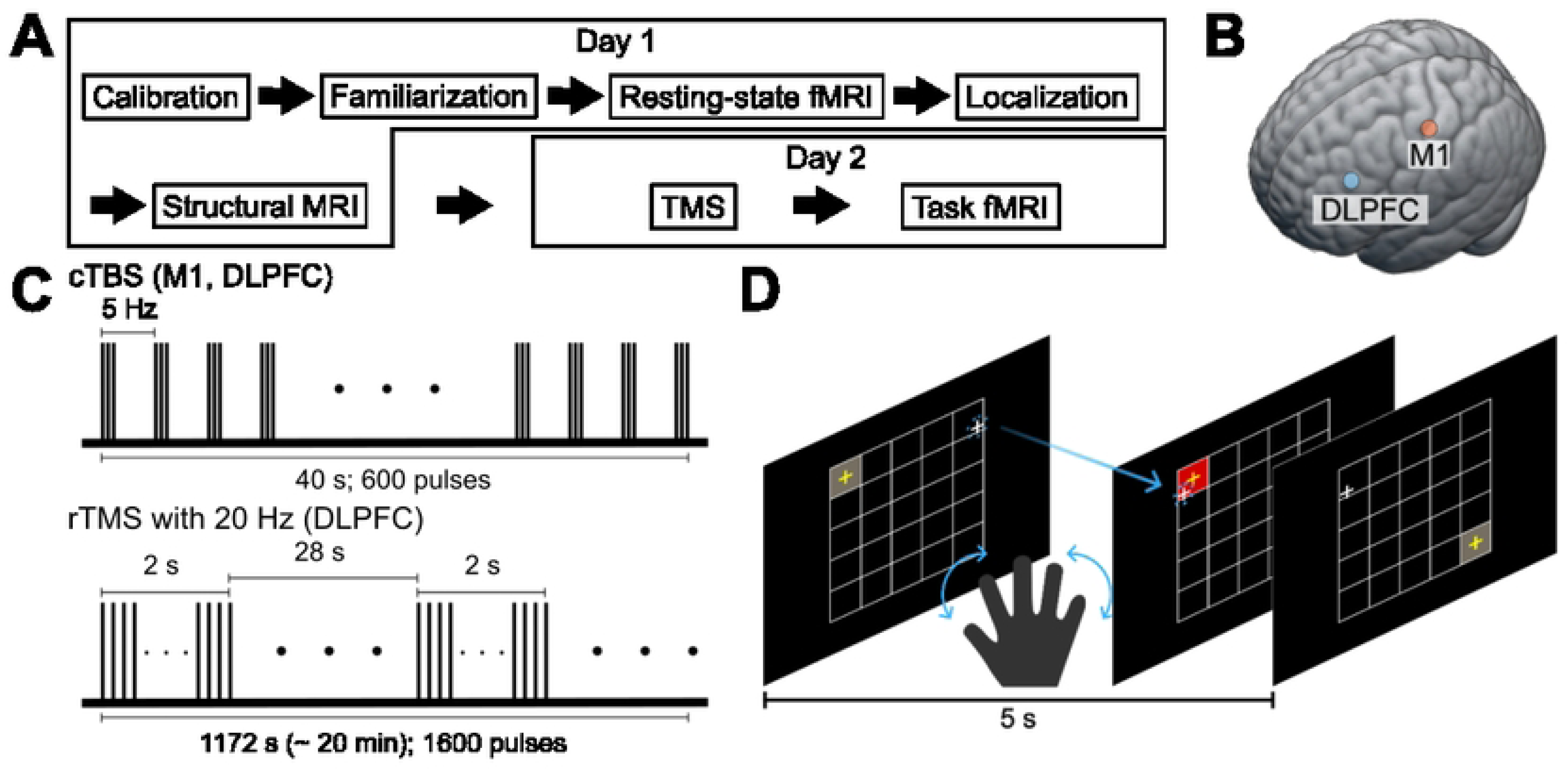
Experimental design and task overview. (A) Experiment schedule. On the first visit, participants underwent familiarization and calibration for behavioral tasks to determine hand-to-cursor mapping. This was followed by three scanning sessions: a resting-state fMRI session, a localization session to identify brain regions associated with random finger movements, and a structural MRI session. On the second visit, participants received TMS to either M1 or DLPFC, then immediately moved into a scanner to perform the motor learning task. (B) TMS target locations in standard space (LPI coordinates): M1 (MNI -38, -22, 52) and DLPFC (MNI -42, 32, 32). (C) cTBS and 20 Hz stimulation protocols. (D) Overview of the motor learning task. Participants learned to move a cursor (white crosshair) to reach a gray target as quickly as possible within 5 seconds. When a target was reached, it turned red. A hand posture represented by 14-dimensional was linearly mapped onto the two-dimensional position of the cursor.

Targets appeared in the four grid corners following a fixed 12-trial block sequence, with all numbers referring to target IDs: 5 (top-right), 25 (bottom-right), 21 (bottom-left), 1 (top-left), 25, 5, 21, 25, 1, 21, 5, 1. As the first trial of each run started centrally, it was excluded from analysis, yielding 96 trials per run (288 total).

### MRI data acquisition

Functional MRI data were collected using a 3-T Siemens Magnetom Prisma scanner equipped with a 64-channel head coil. To identify TMS target sites, a high-resolution anatomical T1-weighted scan was acquired for each participant during the initial visit. For anatomical reference, a whole-brain T1-weighted image was obtained using a magnetization-prepared rapid acquisition with gradient echo (MPRAGE) sequence with the following parameters: repetition time (TR) = 2,400 ms; echo time (TE) = 2.28 ms; flip angle (FA) = 8°; voxel size = 1.00 x 1.00 x 1.00 mm; and field of view (FOV) = 208 × 256 × 256 mm (R-L; A-P; I-S). During the second visit, participants underwent the main experimental sessions, during which functional imaging data were acquired using an echo planar imaging (EPI) sequence with these settings: 1,096 volumes; TR = 460 ms; TE = 27.2 ms; FA = 44°; voxel size = 2.683 × 2.683 × 2.700 mm; and FOV = 220 × 220 × 151 mm. Prior to the functional scans, two EPI images with reversed phase-encoding directions (posterior-to-anterior and anterior-to-posterior) were obtained to facilitate subsequent distortion correction. The parameters for these scans were as follows: TR = 7,220 ms; TE = 73 ms; FA = 90°; voxel size = 2.683 × 2.683 × 2.700 mm; and FOV = 220 × 220 × 151 mm. This MR scan protocol employed a functional imaging sequence identical to that used in previous studies, from which we adopted data as the baseline condition (No-stim)[8].

### Preprocessing

All functional data were preprocessed with AFNI software (version 21.2.04)[39,40] using the default *afni_proc.py* pipeline. Functional images from both sessions first underwent attenuation of extreme time-point outliers (“spikes”) with *3dDespike*, then slice-timing correction (*3dTshift*) and six-parameter rigid-body head-motion correction (*3dvolreg*). The resulting time-series were co-registered to each subject’s high-resolution T1-weighted anatomical scan, nonlinearly warped to the MNI152 template, and resampled to 2.50 mm isotropic voxels. Finally, volumes were spatially smoothed with a 4 mm FWHM Gaussian kernel and intensity-scaled so that each voxel’s time-series had a mean of 100 and ranged from 0 to 200.

The preprocessing pipeline described above was applied uniformly across all datasets to ensure methodological consistency across studies. This included (i) the functional data collected in the present experiment and (ii) the dataset from Choi et al. (2020), which served as a baseline dataset without TMS.

### Behavioral and fMRI data analysis

#### Behavior data analysis

In this experiment, participants were instructed to maintain the on-screen cursor within the designated target cell, which changed every 5 seconds, for as long as possible during each trial. Accordingly, task performance was quantified as the proportion of time (ranging from 0 to 1) that the cursor remained within the target cell during a given trial, and this measure was defined as the success rate. The mean success rate for each of the four participant groups was then determined using two complementary approaches.

First, success rates were averaged within individual blocks (12 trials per block, 24 blocks in total; Fig 2A). This procedure reduced trial-by-trial variability and allowed us to examine improvements in task performance across the course of the experiment for each group. The resulting performance trajectories were approximated using a linear function, and further details of this analysis are provided in the description of the linear mixed-effects model (LME).

**Fig 2.**
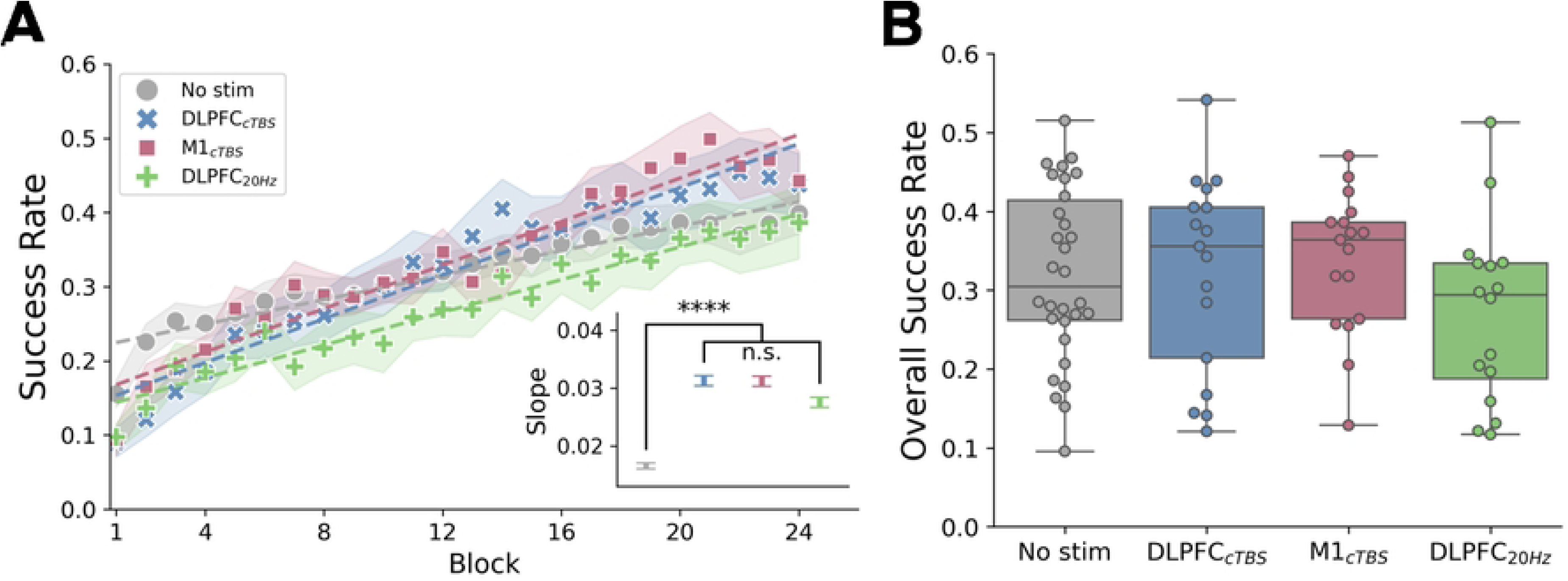
Learning curves and performance differences across groups. (A) The experiment comprised three runs, each consisting of eight blocks (24 blocks in total). Participants’ performance (success rate) was calculated as the average performance per block. Each dot represents the group-level mean, with shaded areas denoting the standard error of the fixed effect regression coefficients. Dashed lines indicate the learning fit for each group as determined by the linear mixed-effects model. The inset graphs illustrate the learning slope for each group. The asterisk above indicates a statistically significant difference between the No-stim group (gray) and the other groups, whereas no differences were observed when comparing only the TMS-stimulation groups. Error bars represent the standard error of the fitted fixed effects coefficients. (B) Box graphs depict the overall performance of participants across the four experimental conditions. No significant differences were observed among groups.

Second, the overall mean success rate was calculated for each participant across all trials (Fig 2B). This measure was used to assess group-level differences in average task performance, for which a one-way ANOVA was conducted.

### fMRI data analysis

Following preprocessing, a univariate voxel-wise general linear model (GLM) was estimated for each participant and run [41]. Trial-by-trial performance, defined as the success rate, was modeled as a parametric modulator, generated by convolving a pulse regressor placed at trial end with a canonical hemodynamic response function [8,27]. Performance, defined as the proportion of time the participant’s cursor remained within the target cell, was conceptualized as an intrinsic reward delivered 5 s after trial onset. The subject-level design matrix included this parametric regressor, six rigid-body motion parameters to account for head movement, and five polynomial regressors (constant plus first through fourth order) to model low-frequency drifts.

To assess the time-dependent effects of TMS on striatal activation, we fitted LME to the GLM-derived parameter estimates extracted from eight predefined striatal subregions (anterior and posterior caudate and putamen in both hemispheres) across the three runs (approximately 10, 20, and 30 minutes after stimulation). Analyses were carried out in Python (version 3.9) using the *statsmodels* package (version 0.14.4). The fixed-effects structure included stimulation condition (DLPFC-cTBS, M1-cTBS, DLPFC-20Hz, and No-stim), run (time), and their interaction, while a random intercept was specified for each participant to account for inter-individual variability in activation patterns.

In addition to model-based estimation, a two-way analysis of variance (ANOVA) was applied to assess the influence of stimulation group and region of interest (ROI) on striatal activity measured at the initial and final runs. This allowed assessment of ROI-specific temporal effects and potential differences in the rate of neural modulation across stimulation conditions. When significant main effects or interactions were identified, we computed the change in evoked activation (30 min - 10 min) for each ROI and conducted post-hoc pairwise comparisons between each stimulation group and the baseline (No-stim) group using Bonferroni-corrected two-tailed t-tests.

To quantify the magnitude of the stimulation-induced modulation, effect sizes were calculated as partial eta-squared (η²). The threshold for statistical significance was set at p < 0.05 (Bonferroni-corrected for multiple comparisons). All analyses were executed using *statsmodels*, and visualization of ROI-wise effects was performed with *matplotlib* and *seaborn*.

### Defining sub-ROIs of the striatum for fMRI analysis

ROIs for the caudate and putamen were generated by identifying overlapping regions between two established neuroanatomical atlases. The first atlas was the TTatlas, specifically the Talairach Daemon database (TTatlas_2010+tlrc.nii)[42], available in AFNI (version 21.2.04). The second atlas was the Harvard-Oxford cortical and subcortical structural atlases, distributed with FSL [43].

For the TTatlas, we used AFNI’s *3dcalc* utility to extract the left and right caudate nuclei (labels: 125, 126, 325, 326) and the putamen (labels: 151, 351) by specifying their corresponding label values. The resulting striatal ROIs were resampled to match the resolution and spatial dimensions of our group-level mask using AFNI’s 3dresample tool.

To confirm the anatomical accuracy of these regions, an independent extraction was performed using the Harvard-Oxford atlas. The left and right caudate (labels: 5, 16) and putamen (labels: 6, 17) masks were created and subsequently resampled into the same group mask space as above.

The final analysis employed the intersection of the caudate and putamen regions across both atlases, ensuring high anatomical fidelity of the ROIs. The caudate ROI was further subdivided into anterior and posterior portions as defined by the TTatlas, which already incorporates this segmentation. For the putamen, anterior and posterior subregions were defined by partitioning at the y = 0 MNI plane, with a one-voxel gap excluded to clearly separate the two halves.

### Linear Mixed-Effects (LME) Models

All LME analyses were implemented in Python (*statsmodels.formula.api*, version 0.14.2) using the *smf.mixedlm()* function. A random-intercept model was specified, with individual subjects included as random effects to account for repeated measurements within participants. We standardized all continuous predictors using z scores prior to fitting the LME models. Standardization improves model stability by reducing scale differences across predictors, which is particularly important for mixed effects optimization procedures that rely on well-conditioned variance–covariance estimates. Z scoring also facilitates the interpretation of fixed effect coefficients, allowing them to be understood as the expected change in the outcome associated with a one–standard deviation increase in each predictor. For these reasons, z score standardization is widely recommended in LME analyses and was applied consistently across all models in the present study. The statistical significance was set at p < 0.05.

#### Model 1-Behavior: Effect of Time (i.e., Block) on Success Rate

We first tested whether time predicted changes in performance within the no-stimulation (No-stim) group:

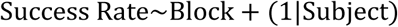

Here, the intercept (‘1’) represents the initial success rate at time = 0 in the reference group (No-stim), serving as the model’s starting point, while the fixed effect of time reflects the slope of change in performance across repeated time points.

#### Model 2-Behavior: Block × Group Interaction

To examine the effects of stimulation and its interaction with block, we fitted the following model:

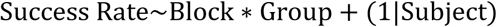

In this specification, the No-stim group served as the reference. This model tested both the main effects of block and group, and critically, whether trajectories of performance over blocks differed between groups.

#### Model 3-fMRI: Effect of Time (i.e., Run) for *β* values

To assess changes in evoked striatal activation (*β* values) during learning in the absence of stimulation (No-stim), we fitted:

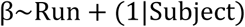

The intercept (‘1’) represents the initial value of the dependent variable (*β*) at time = 0 in the reference group (No-stim), serving as the model’s starting point, while the fixed effect of time reflects the slope of change in *β* across learning, indicating how striatal activation varies as time progresses.

#### Model 4-fMRI: Run × Group Interaction

To evaluate stimulation effects on evoked striatal activity (*β* values), we fitted:

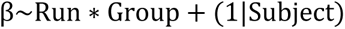

In the model, the fixed effect of time (i.e., Run) captures run-wise changes in striatal activation, the group factor tests overall differences in *β* values between stimulation and No-stim conditions, and the run × group interaction assesses whether the trajectory of striatal activation across runs differs between groups, indicating stimulation-related modulation of learning.

## Results

Fifty participants completed the two-day experiments consisting of two fMRI sessions and one TMS session. Behavioural and MRI data were collected over two days, with the first day involving T1 image acquisition to identify a stimulation site and calibration of the data-glove used in the main experiment for an individual participant. On the second day, participants received TMS immediately followed by a motor learning task during three fMRI runs (Fig 1A), each of which lasted approximately 10 minutes. They performed the motor task in which they wore a data-glove on their right hand and controlled an on-screen cursor by moving their right fingers to reach a target appeared on a screen. The cursor position was calculated from predefined hand-to-cursor mapping, which was calibrated in the first visit for individual participants (see Methods). Participants were randomly assigned to three TMS conditions, cTBS on DLPFC (DLPFC-cTBS) and M1 (M1-cTBS), and 20-Hz rTMS on DLPFC (DLPFC-20Hz) (Fig 1B and 1C). We used cTBS on different cortical regions, DLPFC and M1, as an active control group to each other instead of using a sham control group to investigate the region-specificity of the stimulation effects. Additionally, we also adapted data of 30 participants without TMS as a baseline condition (No-Stim) from our previous study using the same task [8]. The lower learning rates in No-stim group compared to the three stimulation groups was mainly due to the initially greater performance (LME analysis; DLPFC-cTBS: intercept difference = −0.071, p < 0.042; M1-cTBS: intercept difference = −0.057, p < 0.10; DLPFC-20Hz: intercept difference = −0.081, p < 0.023). Participants in No-Stim group were likely to perform the task better initially without a prior TMS procedure. This could be a non-specific effect that was not controlled in our experiment.

Analyses of group-specific learning rates, estimated as slopes using an LME model (see Methods), demonstrated that the cTBS groups exhibited significantly faster performance improvement relative to the No-stim group (i.e., higher learning rates) (Fig 2A). Specifically, the DLPFC-cTBS group showed a z score of 7.55 (Bonferroni-corrected p < 0.0001), and the M1-cTBS group exhibited a z score of 7.46 (corrected p < 0.0001). The DLPFC-20Hz group also showed a modest but significant effect (z score = 3.17, corrected p = 0.0137) (Fig 2A). However, the overall performance among the four groups was not different (Fig 2B; ANOVA, F(3,76) = 1.11, p = 0.35, η² = 0.042).

To investigate the time-dependent modulatory effects of different stimulation protocols on task-related fMRI activity during motor skill learning, we assessed performance-related responses across three fMRI runs conducted approximately 10, 20, and 30 minutes following TMS (Fig 1C). We specifically examined how TMS applied to the DLPFC and M1 selectively modulates fMRI responses within striatal subregions (Fig S2). Based on the known functional connectivity between the anterior and posterior striatum and these cortical areas [18,44], we hypothesized that cTBS targeting the DLPFC and M1 would respectively downregulate activity in the anterior and posterior striatum due to the well-established inhibitory effects of cTBS. To test this hypothesis, we defined regions of interest (ROIs) encompassing eight striatal subregions: anterior and posterior divisions of the caudate and putamen for both hemispheres (Fig S1A). Using voxel-wise general linear model (GLM) analysis with a parametric regressor modeling time-varying performance every second (see Methods), we confirmed prior findings by identifying highly specific task-related responses in the striatum and VMPFC [27].

Furthermore, an LME analysis revealed that, in the baseline group without TMS, the evoked activity across all ROIs decreased significantly over time (i.e., run) (with the smallest absolute z-value observed in the LaCA region; z = –2.872, corrected p = 0.004; Fig S1B). Comparisons between each stimulation group including the baseline group indicated no statistically significant differences during the first 10 minutes post-stimulation, although the effect approached significance and suggested a trend (groupROI two-way ANOVA: F(3,608) = 2.56, p = 0.054, η² = 0.012). In contrast, at 30 minutes post-stimulation, group differences became pronounced (F(3,608) = 12.43, p < 0.0001, η² = 0.058). Moreover, at this time, significant differences across ROIs were also observed (F(7,608) = 9.57, p < 0.0001, η² = 0.099). Subsequently, analysis of evoked activity changes between 10 and 30 minutes demonstrated that, relative to the baseline No-stim group, only the DLPFC-cTBS group exhibited a significant reduction in the left anterior caudate—the region ipsilateral to stimulation (T(45) = 3.22, corrected p = 0.019; Fig 3, Table 1)—indicating a specific inhibitory effect. Similar, but less pronounced reductions were observed in other caudate subregions, whereas no significant modulation was observed in the putamen or following M1-cTBS and DLPFC-20Hz stimulation (Fig 3, Table 1). Collectively, these findings support that network-targeted cTBS modulates corticostriatal circuits in a regionally selective and time-dependent manner during motor learning.

**Fig 3.**
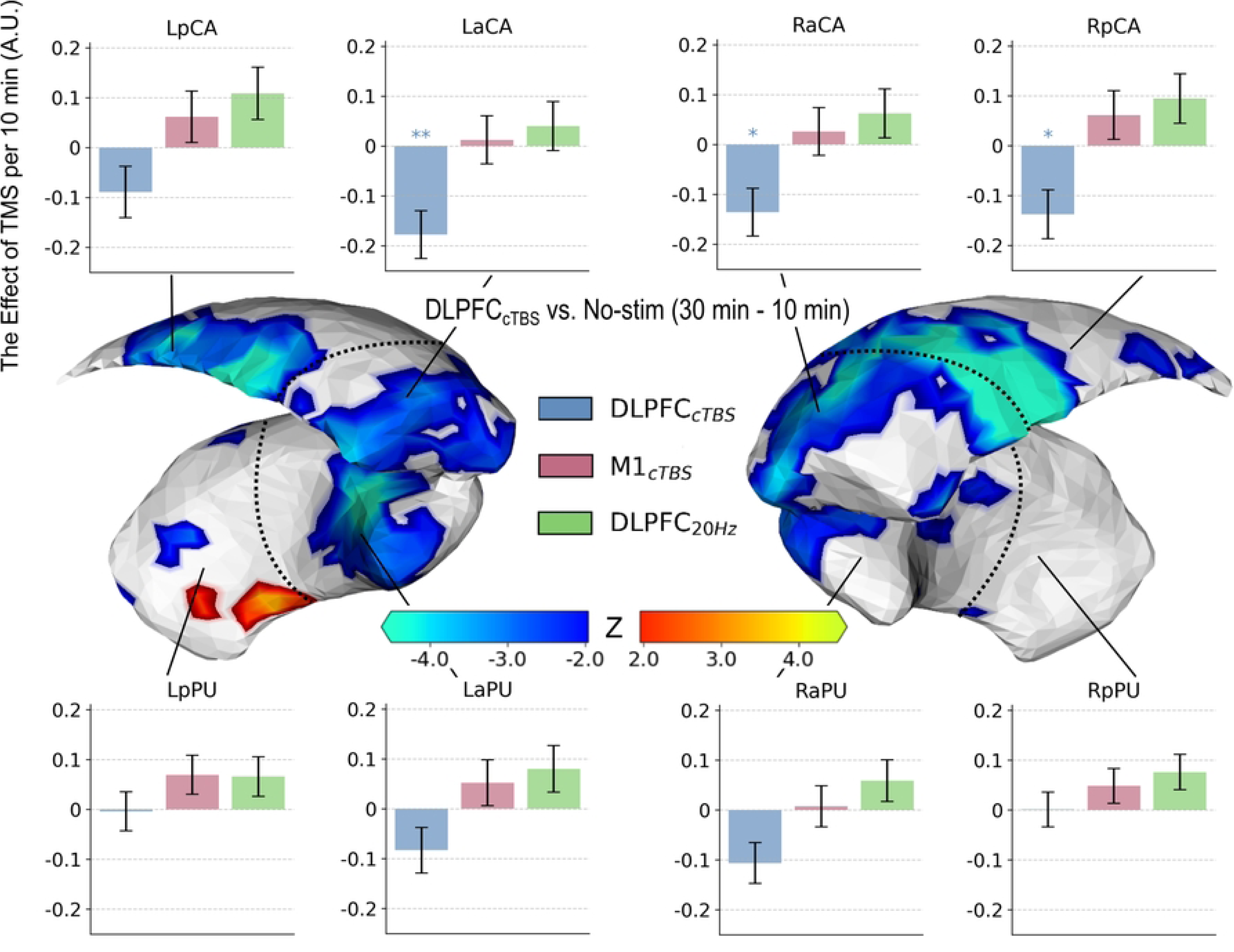
Striatal performance-modulated activation differences between groups over time. Differences in performance-modulated activation between the baseline group and the DLPFC-cTBS group over time in the left and right striatum. (Middle) The rendered image of the caudate nucleus and putamen illustrates regions with performance-modulated activity changes, with the color scale representing Z scores (blue to red: -4.5 to 4.5; the gap in the middle indicates non-significant regions, p ≥ 0.05). (Top and Bottom) Each graph represents the LME results showing differences in activity changes between groups compared to the baseline for each sub-ROI of the striatum. Here, p-values are corrected multiple tests for eight ROIs, and asterisks indicate the following: *, p < 0.05; **, p < 0.01; ***, p < 0.001; ****, p < 0.0001. Error bars represent the standard error of the regression estimates across participants, indicating variability in model-derived activation changes.

**Table 1.**
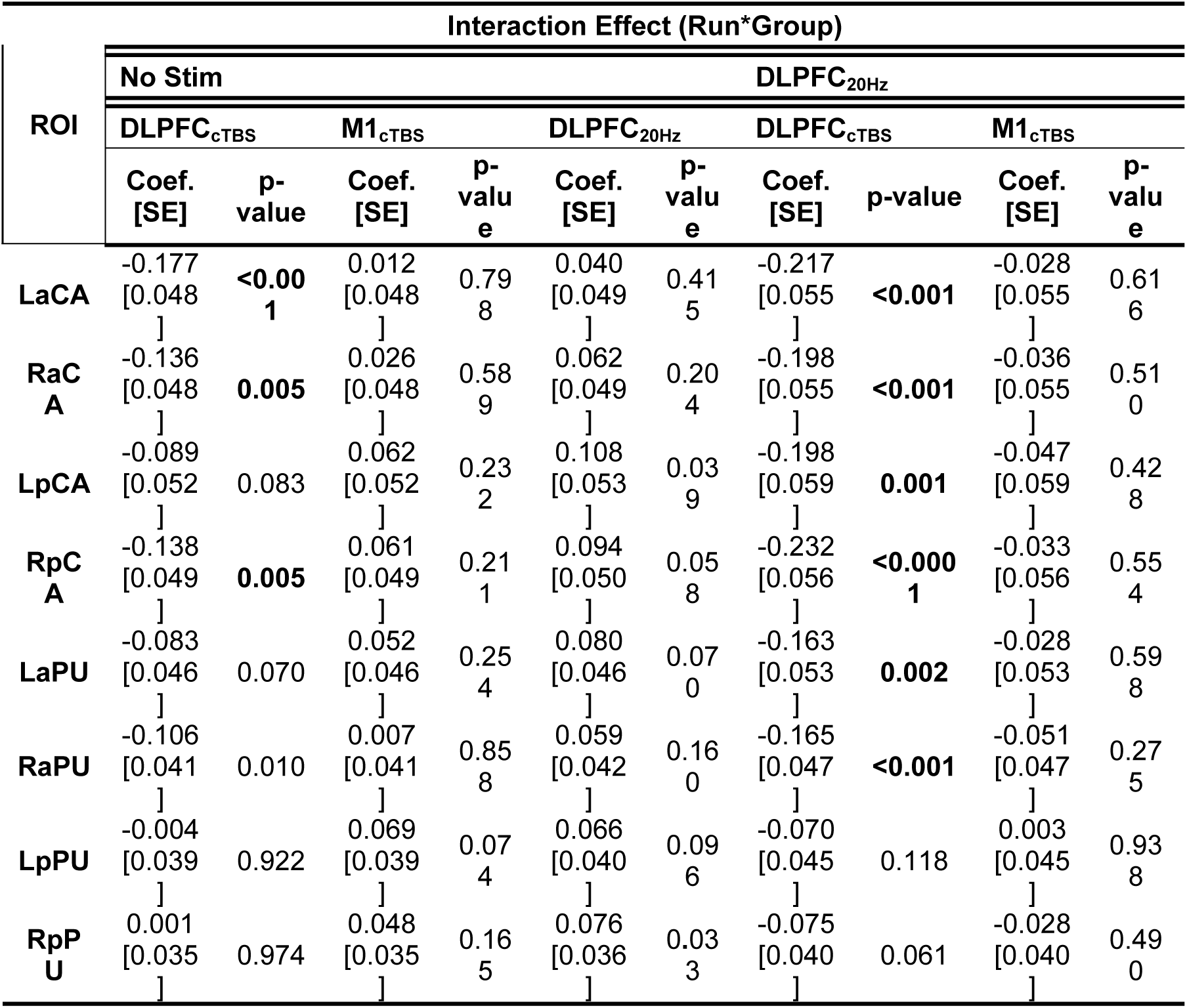
The LME results indicate that the differences between the two groups are attributable to the effect of time (i.e., Run). The *p*-values presented in bold remain statistically significant (*p* < 0.05) even after Bonferroni correction.

To further analyze time-dependent stimulation effects, we assessed the stimulation effect as a relative change in the evoked activity due to stimulation compared to the baseline. Then, we tested whether DLPFC-cTBS modulated the evoked activity in the anterior caudate as a previous study found targeting the exact same site in DLPFC with TMS induced dopamine release in the region [18]. Indeed, we found the most significant effect of DLPFC-cTBS in the left (i.e., ipsilateral to the stimulation site) anterior caudate (T(45) = 3.22 corrected p = 0.019) among eight striatal subregions, which was greater reduction in the evoked activity than the baseline (Fig 3, Table 1). In this region, DLPFC-cTBS group showed significantly greater stimulation effect than other TMS groups (vs M1-cTBS: T(32) = 2.94, corrected p < 0.05, vs DLPFC-20Hz: T(31) = 3.69, corrected p < 0.01), where no stimulation effect (compared to the baseline) was observed (M1-cTBS: T(45) = 0.24, DLPFC-20Hz: T(44) = 0.85) (Fig 3, Table 1). Thus, the downregulation of the evoked activity in the left anterior caudate was specific to DLPFC region and DLPFC-20Hz did not induce any excitatory effects on the evoked activity in the region. A similar effect due to DLPFC-cTBS, although less pronounced, was found in other striatal subregions such as the right anterior (T(45) = 2.55, corrected p = 0.11) and the right posterior caudates (T(45) = 2.37, corrected p = 0.18). Interestingly, the inhibitory effect of DLPFC-cTBS was specific to the caudates, not in the putamen (Fig 3).

Finally, we also tested whether M1-cTBS modulated the evoked activity in the posterior striatal regions. Previous studies demonstrated that targeting the right M1 at 1 Hz (inhibitory)[20] or 10 Hz (excitatory)[44] induced weaker between the M1 and posterior putamen functional connectivity and elicited distinct patterns of dopamine release in the same region. However, we did not find a significant stimulation effect in any striatal subregions. Consistently, DLPFC-20Hz stimulation did not modulate the evoked activity in any striatal subregions (Fig 3, Table 1).

## Discussion

Motor learning is known to rely on the dynamic interaction between cortical and subcortical circuits, especially the corticostriatal network. In this study, we investigated the role of the corticostriatal network in motor learning and how it contributes to task acquisition using TMS. As a result, we found that continuous theta-burst stimulation of the dorsolateral prefrontal cortex (DLPFC-cTBS) reduced task-related striatal fMRI activity, although there was no significant difference in overall behavioral performance between stimulation groups. One plausible explanation for the lack of behavioral effects involves individual differences in baseline strategy preference, working memory capacity, and reward sensitivity. Large-scale studies have demonstrated considerable inter-individual variability in behavioral and neural responses to theta-burst stimulation [45]. Such variability may mask group-level behavioral changes even if consistent neural modulations, such as reductions in striatal BOLD signals, are present.

Previous research using the same task has shown that time-varying task performance modulates striatal activity with high sensitivity and specificity, processes closely linked to reward-based learning [8,27]. Notably, these effects were most pronounced in the left anterior caudate nucleus, a region anatomically connected ipsilaterally to the stimulated left DLPFC, comprising part of the anterior cognitive corticostriatal loop [5,6,8,18]. In our study, stimulation effects were not significant at 10 minutes post-stimulation but emerged after 20 minutes and peaked at 30 minutes, consistent with earlier findings indicating maximal inhibitory cTBS effects on motor evoked potentials between 15 and 45 minutes post-stimulation [15]. However, due to our task’s limited duration (30 minutes), we did not observe the eventual offset of this effect.

In contrast to DLPFC stimulation, cTBS applied over M1 did not modify task-related striatal activity despite previous evidence that 10 Hz rTMS on M1 induces dopamine release in the putamen [44]. This discrepancy may be due to the predominance of anterior cognitive corticostriatal loops during early learning phases of our short task [5,6,8,18]. Therefore, M1-targeted stimulation of the posterior sensorimotor loop may exert more notable effects in later learning stages involving motor memory consolidation, as supported by previous reports showing 1 Hz rTMS disrupts consolidation by reducing connectivity between M1 and dorsal striatum [19–21].

The high frequency 20-Hz rTMS on DLPFC, which we hypothesized to have an excitatory effect [26,33–35,46,47], also did not induce significant difference in the striatal activity compared to the baseline. Previous studies demonstrated that the 20-Hz rTMS applied on the left lateral parietal region, which is functionally connected to the hippocampus, have excitatory effects in the downstream regions in the posterior-medial corticohippocampal network [26,33–35]. While the effects of 20-Hz rTMS were reproducible in the corticohippocampal network in the multiple studies, they may not the case in the corticostriatal network as shown in our study. However, there still remains a possibility that other excitatory stimulation protocols such as 10-Hz rTMS and intermittent theta-burst stimulation (iTBS) on DLPFC could increase the task-related striatal activity [18,44,48]. In future work, our network-targeted approach may combine these excitatory stimulation protocols to treat patients with Parkinson’s disease, which is characterized by striatal hypoactivity and corticostriatal network dysregulation [17,49].

Multimodal imaging studies combining fMRI and PET have demonstrated close associations between striatal BOLD responses and dopaminergic activity during reward processing [50–52]. Furthermore, human studies using 10-Hz rTMS applied on DLPFC and M1 induced dopamine release ipsilaterally in the anterior caudate and dorsal putamen, which were measured by reduction in [11C]raclopride binding in PET scans [18,44]. Thus, decreased anterior striatum fMRI activity observed after DLPFC-cTBS may reflect dopaminergic downregulation, suggesting potential therapeutic benefits for dopamine-related disorders such as addiction. However, decreased BOLD signals do not necessarily indicate local neural inhibition; studies have shown that outside primary sensory/motor cortices, TMS-fMRI responses often reflect network-level indirect effects [53]. Our results may similarly be mediated by changes in prefrontal-striatal functional connectivity, warranting further TMS-fMRI connectivity analyses.

While our results support the effect of network-targeted TMS on the striatal activity, the underlying mechanism of the effect is still unclear. A recent study employing a concurrent TMS-fMRI has demonstrated that TMS pulses on the ventrolateral PFC triggered significant activity changes in the amygdala [54]. The stimulation effect was more significant with stronger anatomical connectivity between VLPFC and amygdala, revealed by diffusion tensor imaging. This also could be the case with the corticostriatal network and a similar study using TMS-fMRI warrants to provide more mechanistic understanding for our results [14]. Additionally, a longitudinal study with multiple TMS sessions is necessary to investigate induced neural plastic changes such as long-term potentiation (LTP) or depression (LTD) in the corticostriatal network. Finally, temporal interference stimulation (TIS), which is an emerging new noninvasive technique, has been recently applied to directly stimulate the human striatum and enhance motor learning as well as task-related striatal activity [55]. While TIS is considered the only technique to stimulate subcortical regions noninvasively, the effect is still subthreshold, like typical noninvasive electrical stimulation methods. Combining the network-targeted TMS with TIS could be a promising approach to overcome the limitation as TMS may serve as a pre-conditioning stimulation before TIS or vice versa.

In sum, our study demonstrates the validity of network-targeted TMS via combined TMS-fMRI and reveals temporally specific inhibitory effects of cTBS on corticostriatal activity during motor learning, advancing understanding of corticostriatal circuits’ causal roles and neuromodulation’s therapeutic potential for neurological and psychiatric disorders.

## Author Contributions

S. Park and S. Kim designed the research, interpreted the results. J. Kim and Y. Kwon recruited the participants and conducted all experimental sessions. S. Park wrote the paper and analyzed the data and prepared all figures. All authors reviewed the manuscript.

## Acknowledgements

Neuroimaging was performed at the Center for Neuroscience Imaging Research located at Sungkyunkwan University, supported by the Institute for Basic Science.

## Data and code availability

All MRI data used in this study were archived in the Hanyang University Network Attached Storage (NAS). All the dataset and codes related to this study will be publicly available upon the acceptance of the manuscript.

## Supporting information

**S1 Fig. The division of the striatum into eight ROIs and their analysis.** (A) The caudate and putamen, parts of the striatum, are each divided into four regions: anterior (a), posterior (p), left (L), and right (R) for analysis. (B) Results of the mixed linear regression analysis for time-dependent changes in performance-modulated activity in the ‘No-Stim’ group for each ROI. Here, p-values are corrected for ROI comparisons, and asterisks indicate the following: *, p < 0.05; **, p < 0.01; ***, p < 0.001; ****, p < 0.0001.

**S2 Fig. Time-dependent changes in performance-modulated activity across ROIs for each group.** No significant group or ROI differences were observed during the early post-stimulation period (10 minutes). In contrast, group differences emerged during the late post-stimulation period (30 minutes), with the largest divergence observed between the DLPFC-cTBS and DLPFC-20Hz groups. P-values are uncorrected, with significance indicated as follows: *, *p* < 0.05; **, *p* < 0.01; ***, *p* < 0.001; ****, *p* < 0.0001. Error bars represent the standard error of the mean (SEM).

**S3 Fig. Time-dependent whole-brain voxel-wise GLM results showing performance-modulated BOLD activity.** Voxel-wise general linear model analyses were conducted for each TMS group to identify brain regions in which trial-by-trial performance modulated BOLD responses across the approximately 10-minute post-stimulation period. Statistical maps depict regions exhibiting significant performance-related modulation, with color scales indicating Z scores. All maps were thresholded at the group-level criterion of |Z| > 1.96.

